# Nonlocal Proliferation and Explosive Tumour Dynamics: Mechanistic Modelling and Bayesian Inference

**DOI:** 10.64898/2026.03.23.713731

**Authors:** Nikos I. Kavallaris, Farrukh Javed

**Affiliations:** Department of Mathematics and Computer Science, Karlstad University, Karlstad, Sweden; Department of Statistics,Lund University, Lund, Sweden

**Author notes:** Contributing authors. These authors contributed equally to this work.

**Keywords:** Nonlocal reaction–diffusion, convolution kernel, Kawarada quenching, finite-time singularities, Neumann boundary conditions, tumour growth, Bayesian inference, uncertainty quantification

## Abstract

We introduce a mechanistic, nonlocal tumour-growth model designed specifically to capture explosive dynamics that are not adequately explained by standard logistic reaction–diffusion descriptions. The motivation is empirical: the universal scaling law reported in [1] provides compelling cross-sectional evidence of superlinear tumour activity versus tumour burden, but as a phenomenological relationship it does not by itself supply a dynamical mechanism, nor does it rigorously describe how explosive growth emerges, how fast it develops, or how spatial interactions and tissue boundaries influence it. Our model addresses this gap by incorporating nonlocal proliferative feedback—cells respond to a spatially aggregated neighbourhood signal—and a singular, Kawarada-type acceleration that produces “quenching”: tumour density stays bounded while the proliferative drive becomes unbounded as the aggregated signal approaches a critical threshold. This offers a concrete mechanistic route to explosive escalation consistent with physical boundedness.

We analyse the model under no-flux (Neumann) boundary conditions, appropriate for reflecting tissue interfaces. In the spatially homogeneous setting we prove finite-time onset of the explosive regime and obtain explicit rates for how rapidly it is approached. For spatially heterogeneous perturbations we derive a transparent spectral stability theory showing how the interaction kernel selects spatial scales and how the singular acceleration tightens stability margins as the explosive threshold is approached. These results provide interpretable links between nonlocal interaction structure, boundary effects, and the emergence of rapid growth.

Finally, to connect mechanism to data in the spirit of [1], we embed the model in a Bayesian inference framework that treats the interaction kernel and the acceleration strength as unknown and learned from tumour-growth observations. This enables uncertainty-aware estimation of explosive onset times, escalation rates, and stability margins, while positioning the scaling law of [1] as an observable signature that our mechanistic model can explain and quantify rather than merely fit.

## 1 Introduction

Classical logistic models are widely used to describe population dynamics in biology, ecology, and medicine, and can be formulated as reaction–diffusion equations with local source terms of Fisher–KPP type [2]. However, many biological systems exhibit feedbacks that depend not only on the local density *u*(*x, t*) but also on the *global* or *mesoscopic* size of the population, typically represented by some spatially averaged or smoothed density. Nonlocal interaction and sensing mechanisms of this kind have been extensively studied in collective dynamics, chemotaxis, and kinetic transport models [3, 4].

The present work is motivated by experimental evidence that human tumours can exhibit *explosive growth* driven by superlinear metabolic scaling laws [1, 5]. Using large multi-centre PET and MRI datasets, Pérez-García et al. [1] demonstrated that key quantitative observables of tumour activity obey power-law relationships with exponents significantly larger than one, indicating strong nonlinear coupling between tumour size and effective metabolic or proliferative activity. Related analyses and stochastic growth models have further connected explosive tumour dynamics to branching processes, metastasis formation, and spatial heterogeneity [6–9].

From a mathematical point of view, our model combines a nonlocal Fisher–KPP-type structure with a singular feedback law. Singular reaction terms that blow up as the solution approaches a threshold have a long history, starting with the work of Kawarada [10], and have been analyzed in a variety of local PDE settings [11–13]. Here we adapt this framework to a convolution-based nonlocal setting inspired by tumour growth, and we focus on homogeneous Neumann boundary conditions, which correspond to reflecting boundaries with no flux of cells through the boundary.

In parallel, advances in Bayesian statistics and computational methods have made it possible to calibrate complex mechanistic PDE models from noisy experimental data, including via likelihood-free techniques such as approximate Bayesian computation [14, 15]. This motivates the Bayesian inference and uncertainty quantification part of our study, in which we aim to learn the interaction kernel and singularity exponent from data in an uncertainty-aware fashion.

### Outline of the paper

The remainder of the paper is organized as follows. In Section 2 we introduce the nonlocal FKPP–Kawarada-type model as a mechanistic description of tumour cell-density evolution in which proliferation is regulated by a spatially aggregated neighbourhood signal encoded by the interaction kernel, and we formulate the corresponding Neumann (no-flux) boundary-value problem to represent reflecting tissue interfaces with no net cell flux across the boundary. Section 3 studies the resulting dynamics, including well-posedness and positivity, finite-time quenching (with explicit quenching-time and quenching-rate results in the spatially homogeneous reduction), and the stability and travelling-wave analysis that elucidates how nonlocal interactions shape spatial structure. Section 4 presents the Bayesian calibration framework and shows how uncertainty in the kernel and singularity exponent propagates to uncertainty in quenching times, quenching rates, and stability margins. We then present the cohort data and statistical workflow in Section 5, and close with limitations and directions for future work in Section 6.

## 2 Model formulation and motivation

### 2.1 The nonlocal model

Recent work in [1] proposed a phenomenological nonlocal model to explain explosive tumour growth observed in clinical imaging data. In that framework, a global tumour activity variable (often interpreted as a total tumour “velocity” or metabolic rate) is coupled to the tumour size through a nonlocal feedback law. At the level of an associated ODE for this aggregate quantity, the authors derive in the Supplementary Material an *upper* solution that can blow up in finite time, thereby providing a mechanism for rapid acceleration of tumour activity. However, the structure of the nonlocal feedback in [1] does not permit a corresponding *lower* blow-up estimate, and, in particular, it does not yield a rigorous blow-up of the time derivative of the solution for the full nonlocal PDE. Explosive behaviour appears as a property of a specific observable (an upper solution of the ODE for total activity), rather than as a robust feature of the underlying spatial dynamics.

The aim of the present work is to introduce a genuinely *singular* nonlocal model, in which the singularity is encoded directly at the level of a proliferation rate depending on a spatial convolution of the tumour density. In this way, explosive tumour behaviour is no longer an artefact of a particular upper solution, but a built-in mechanism of the nonlocal reaction–diffusion dynamics: the reaction term itself blows up as a nonlocal density approaches a critical threshold, while the density remains bounded. At the same time, the model remains compatible with the data-driven calibration philosophy of [1, 5], through a small set of biophysically interpretable parameters such as the strength and range of nonlocal interactions and a singularity exponent.

From a biological point of view, we focus here on *homogeneous Neumann boundary conditions*, which correspond to a no-flux condition for tumour cells across the boundary. This choice is standard in continuum models of tumour invasion and growth when the computational domain represents a portion of tissue that is effectively closed to cell exchange on the time scales of interest; see, for instance, reaction–diffusion and chemotaxis-type models of avascular and vascular tumours in [16–18] and glioma growth models in which the brain or a brain subregion is treated as a reflection domain with no cell flux across artificial boundaries or skull interfaces [19–21]. In such settings, the domain Ω may represent (i) a subregion deep inside a larger organ (so that the boundary of Ω is artificial and should not create spurious loss or gain of cells), or (ii) a tissue compartment separated from surrounding structures by interfaces that act as effective reflecting barriers on the relevant time scales.

Let Ω ⊂ ℝ^*d*^ be a bounded domain with smooth boundary ∂Ω. The unknown *u* = *u*(*x, t*) denotes a scaled cell density, normalized so that 0 *< u <* 1. Under homogeneous Neumann boundary conditions the proposed model reads

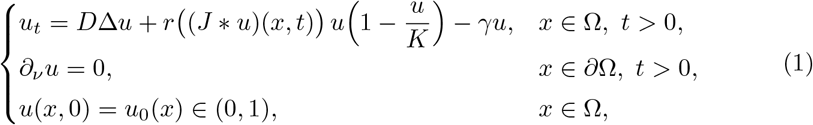

where ∂_*ν*_*u* denotes the normal derivative of *u* in the direction of the outward unit normal vector *ν* on ∂Ω. Biologically,

- *D >* 0 represents effective cell motility (random dispersal in tissue);
- 0 *< K* ≤ 1 is a local carrying capacity summarising microenvironmental limits (space/nutrient/oxygen constraints);
- *γ* ≥ 0 is an effective loss rate (death/clearance or net removal);
- (*J* * *u*)(*x, t*) is a neighbourhood-averaged tumour burden,

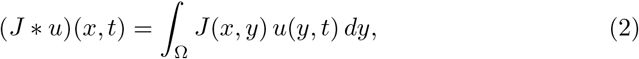

where *J* encodes the spatial range of cell–cell and microenvironmental coupling (e.g. paracrine signalling or mechanical influence; cf. Figure 1);
- *r*( · ) is a nonlocal proliferation rate regulated by the aggregated field *w*(*x, t*) := (*J* * *u*)(*x, t*).

**Figure 1.**
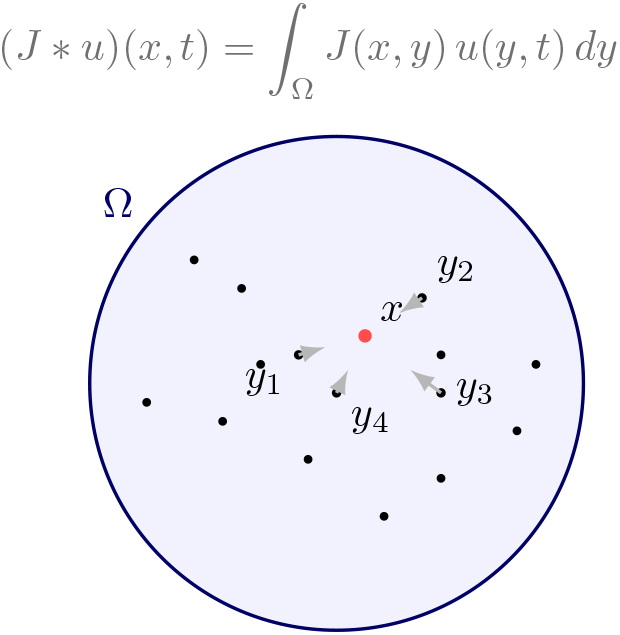
Nonlocal kernel interaction in a circular tumour domain Ω. The effective proliferation at *x* depends on the weighted average (*J* * *u*)(*x, t*) = *∫*_Ω_ *J* (*x, y*) *u*(*y, t*) *dy* over neighbouring cells *y*.

We assume:

(K1) J ∈ C(Ω × Ω), J(x, y) ≥ 0 for all x, y ∈ Ω;

(K2) normalization *∫* _Ω_ *J*(*x, y*) *dy* = 1 for each *x*;

(K3) positivity: there exists *κ >* 0 such that *J*(*x, y*) ≥ *κ* for all *x, y* ∈ Ω.

Guided by nonlocal tumour-growth models [1, 5], we prescribe the nonlocal proliferation rate as

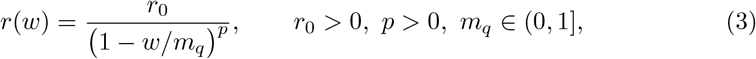

where *m*_*q*_ is a critical nonlocal density at which *r*(*w*) diverges.

Substituting (3) into (1) yields

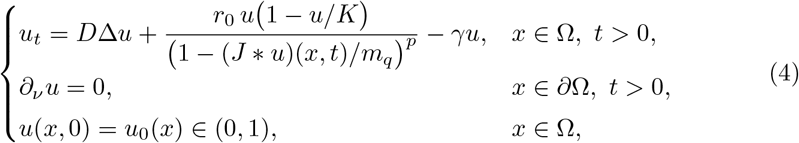

a nonlocal analogue of a Kawarada quenching equation [10, 22–24].

- The singular dependence *r*(*w*) ∝ (1 − *w/m*_*q*_)^−*p*^ captures the idea that proliferation rates accelerate dramatically as the convolved density approaches a critical value *m*_*q*_, which may correspond to a threshold in vascularization or metabolic support beyond which further growth enhances per-cell activity.
- From a broader perspective, the convolution kernel *J* can be viewed as a coarsegrained representation of the vascular and metabolic coupling between different tumour regions, while the singular feedback *r*(*w*) packages complex biophysical processes into an effective law. The analysis presented here thus provides a mathematically transparent framework that can be embedded into more detailed models of cancer growth and treatment response.

In [1] the authors considered a linear nonlocal dependence of the proliferation rate of the form

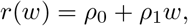

for some positive constants *ρ*_0_, *ρ*_1_, with *γ* = 0 and with the kernel chosen as *J* = *δ*_0_ (Dirac mass at a source point *x* = 0). This leads to the nonlocal proliferation model

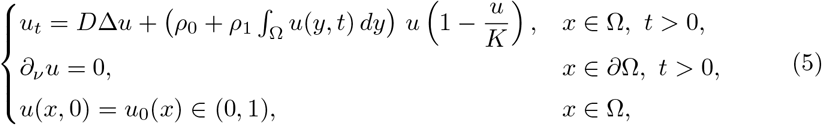

in which the global tumour burden enters only through a linear feedback on the growth rate. While this model can exhibit rapid growth at the level of suitable upper solutions, it does not enforce the genuinely explosive tumour behaviour described above, in the sense that no rigorous quenching or blow-up mechanism can be derived for the full PDE from this linear nonlocal coupling.

Beyond tumour-growth motivations, closely related *nonlocal* quenching-type models have also been studied in the applied-PDE literature on micro-electro-mechanical systems (MEMS), where nonlocal terms arise through electrostatic coupling; see [24– 26]. *This MEMS literature is also directly connected to the present work: the relevant expertise of some of the authors in nonlocal MEMS-type quenching models helped motivate the introduction of the nonlocal Kawarada-type feedback mechanism proposed here for explosive tumour dynamics*. Complementary insight is provided by *stochastic* local counterparts that capture random fluctuations in the reaction dynamics, analysed in [27–30].

### 2.2 Biological and data-driven motivation

The specific choice of a singular, nonlocal proliferation rate

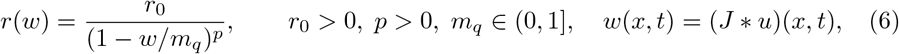

is motivated by the superlinear metabolic scaling laws reported in Pérez-García et al. [1]. In that study, volumetric PET imaging data showed that metabolic activity and proliferation scale with tumour volume *V* as a power law of the form

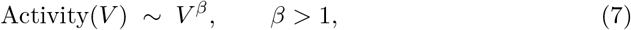

with exponents typically around *β* ≈ 5*/*4, implying that larger tumours have disproportionately higher specific proliferation rates and that the associated continuous growth law develops a finite-time singularity if extrapolated. To rationalise these findings, [1] derived a nonlocal Fisher–KPP model in which the proliferation coefficient depends linearly on the global tumour mass *N* (*t*) = *∫*_Ω_ *u*(*x, t*) *dx*, thereby reproducing the observed superlinear scaling.

More recent longitudinal MRI studies of brain metastases have confirmed and refined this picture. Ocaña-Tienda et al. [5] analysed large cohorts of human brain metastases and showed that untreated lesions are well described by a von Bertalanffy-type law 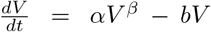 with growth exponents *β >* 1, corresponding to super-exponential volumetric growth, whereas treated lesions typically exhibit *β <* 1. From a stochastic and evolutionary viewpoint, branching-process models of secondary tumours [15] and evolutionary models with random fitness advances [6] predict similar super-exponential or even finite-time blow-up behaviour for tumour cell populations, while allometric models emphasise the connection between tumour metabolism and growth exponents [7]. Collectively, these works support the view that explosive growth in malignant tumours emerges from evolutionary dynamics in a heterogeneous population, coupled to metabolic up-regulation as size increases.

Our law (6) can be viewed as a nonlinear, spatially refined generalisation of the above mechanisms. For small values of the nonlocal tumour load *w*, the expansion

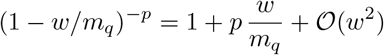

shows that

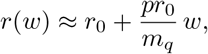

which recovers, to leading order, a size-dependent proliferation rate of the form *r*(*w*) ≈ *ρ*_0_ + *ρ*_1_*w* consistent with the nonlocal Fisher–KPP model (5) proposed in [1]. Here *w*(*x, t*) = (*J * u*)(*x, t*) is a coarse-grained measure of the tumour burden or metabolic environment around *x*, reflecting the fact that evolutionary selection of aggressive clones and metabolic up-regulation act over a mesoscopic neighbourhood rather than at a single point.

As *w*(*x, t*) ↗ *m*_*q*_, the factor (1 − *w/m*_*q*_)^−*p*^ diverges and produces an idealised “explosive” regime in which the proliferation rate accelerates rapidly under nonlocal feedback, mirroring the finite-time singularity that emerges from sustained superlinear scaling. The exponent *p >* 0 controls the strength of this acceleration, in direct analogy with the effective growth exponents *β* estimated from volumetric data [1, 5], while *m*_*q*_ represents a critical nonlocal load beyond which the continuum model predicts runaway growth. In this sense, the nonlocal Kawarada structure of our model provides a mesoscopic PDE interpretation of the empirically observed tumour scaling laws and explosive growth behaviour.

## 3 Model dynamics

Since the nonlocal model (4) is, to the best of our knowledge, new in the context of cancer dynamics, we devote this section to a detailed analysis of its well-posedness and positivity properties, its long-time behaviour and linear stability, and to a travelling-wave study in the one-dimensional setting.

### 3.1 Well-posedness and positivity

We first establish local well-posedness and positivity preservation for the nonlocal Kawarada model with Neumann boundary conditions. Recall that (4) can be written in the form

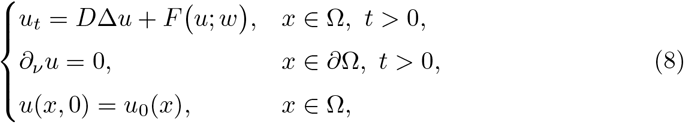

where

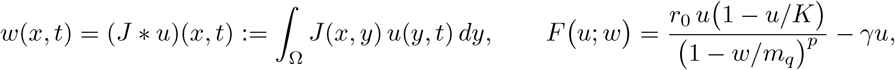

and *J* satisfies (K1)–(K3). The singularity of the reaction term occurs when *w* approaches *m*_*q*_, so we restrict attention to the region

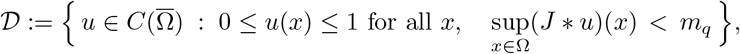

on which the denominator 1 − *w/m*_*q*_ stays uniformly away from zero.

#### Local existence and uniqueness

Let 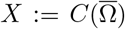 equipped with the sup norm 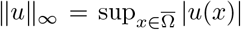. The Neumann Laplacian *A* := *D*Δ with domain

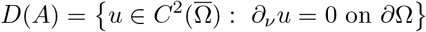

generates an analytic semigroup (*e*^*tA*^)_*t*≥0_ on *X* (see, e.g., [31]). The nonlocal operator *u* → *w* = *J* * *u* is a bounded linear map *X* → *X* under (K1)–(K2), since

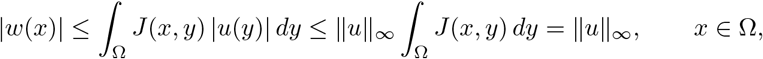

so ∥*J* * *u*∥_∞_ ≤ ∥*u*∥_∞_.

On any bounded subset of *X* satisfying sup_*x*∈Ω_(*J* * *u*)(*x*) ≤ *m*_*q*_ − *η* for some fixed *η >* 0, the map

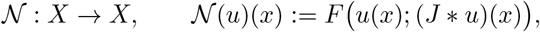

is locally Lipschitz. Indeed, *u* → *J* * *u* is linear and bounded, and (*u, w*) → *F* (*u*; *w*) is smooth on {(*u, w*) ∈ ℝ^2^ : 0 ≤ *u* ≤ 1, *w* ≤ *m*_*q*_ – *η*}. Therefore the nonlocal Kawarada equation (8) is a semilinear parabolic problem with locally Lipschitz nonlinearity on 𝒟. Standard semigroup theory (see, e.g., [31] and references therein) then yields:

##### Proposition 1

(Local well-posedness). *Let* 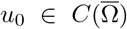 *satisfy* 0 ≤ *u*_0_ ≤ 1 *and* sup_*x*∈Ω_(*J* * *u*_0_)(*x*) *< m*_*q*_. *Then there exists T* ∈ (0, ∞] *and a unique classical solution*

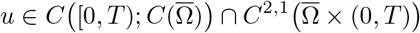

*of* (8) *such that u*(·, 0) = *u*_0_. *Moreover, either T* = ∞, *or T <* ∞ *and*

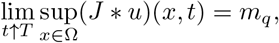

*i.e*., *finite-time approach to the nonlocal singularity is the only obstruction to extending the solution*.

#### Positivity and uniform bounds

We now show that solutions preserve nonnegativity and stay uniformly bounded (as long as they exist). First note that *u* ≡ 0 is a stationary solution of (8), since *F* (0; *w*) ≡ 0. Under (K1)–(K3), the reaction term satisfies

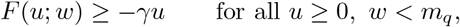

and the convolution *J* * *u* maps nonnegative functions to nonnegative functions. Thus the comparison principle for parabolic equations with Neumann boundary conditions (cf. [31]) implies that if *u*_0_ ≥ 0 then

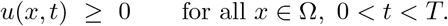

Similarly, the logistic structure bounds *u* from above. Suppose *u* attains a spatial maximum at (*x*_0_, *t*_0_) with *u*(*x*_0_, *t*_0_) *> K*. At such a point one has Δ*u*(*x*_0_, *t*_0_) ≤ 0 and ∂_*ν*_*u* = 0 on ∂Ω, so

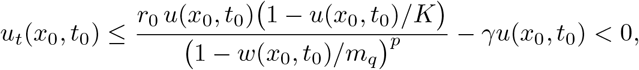

because 1 −*u/K <* 0 when *u > K*. This contradicts the maximality of *u*(*x*_0_, *t*_0_) unless *u*(*x, t*) ≤ *K* for all (*x, t*) in the domain of existence. In particular, if 0 ≤ *u*_0_ ≤ 1 and *K* ≤ 1, then

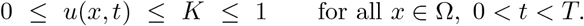

Combining this with the bound ∥*J* * *u*(, *t*) ∥_∞_ ≤ *u* ∥ (, *t*) ∥ _∞_ shows that, as long as *t < T*, the solution remains in the invariant region

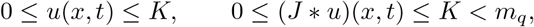

and hence the singularity is not encountered before *T* .

In summary, under our kernel assumptions (K1)–(K3) and for biologically relevant initial data 0 ≤ *u*_0_ ≤ 1, the nonlocal Kawarada model (4) admits a unique classical solution on a maximal time interval [0, *T*_max_), preserves nonnegativity, and keeps the density uniformly bounded by *K* as long as it exists. The only possible loss of well-posedness is through the nonlocal quantity *w* = (*J* * *u*) approaching the critical threshold *m*_*q*_, which corresponds precisely to the quenching scenario analysed in the next section.

### 3.2 Quenching Behaviour

Set

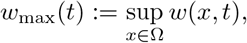

and recall the reaction term

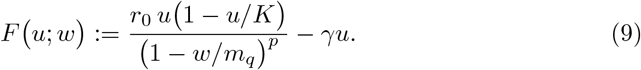

#### Definition 2

(Quenching for the Neumann problem). A classical solution *u* of (4) *quenches* at time *T*_*q*_ *<* ∞ if:

- *u*(*x, t*) ∈ (0, 1) for all *x* ∈ Ω and *t < T*_*q*_, and sup_*x*∈Ω_ *u*(*x, t*) remains bounded on [0, *T*_*q*_);
- 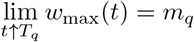
- 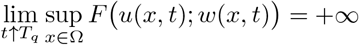

#### Spatially homogeneous reduction

Assume homogeneous Neumann boundary conditions and homogeneous initial data ta *u*0(*x*) = *ū*0 ∈ (0,1). In this case the structure of the equation and the kernel assumptions (K1)–(K2) imply that spatially constant states form an invariant class. Indeed, if we make the ansatz

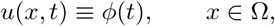

then ∂_*ν*_*u* = 0 on ∂Ω holds automatically and

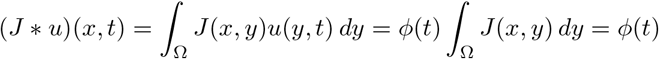

for every *x* ∈ Ω by (K2). Thus the PDE reduces to the scalar ODE for *ϕ*,

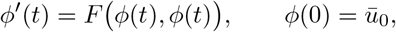

Standard ODE theory yields a unique solution *ϕ*(*t*) on its maximal interval of existence. Defining *v*(*x, t*) := *ϕ*(*t*), we see that *v* satisfies the full PDE with Neumann boundary conditions and initial data *u*_0_ ≡ *ū*_0_. On the other hand, the parabolic problem with these data admits a unique (classical) solution *u* (see, e.g., [31]). Therefore *u* ≡ *v*, i.e.

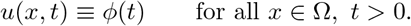

In particular, spatially homogeneous initial data under no-flux boundary conditions generate spatially homogeneous solutions, and the dynamics reduce exactly to the nonlocal Kawarada ODE for *ϕ*. Using assumption (K2), *w*(*x, t*) = (*J* * *u*)(*x, t*) = *ϕ*(*t*), and *w*_max_(*t*) = *ϕ*(*t*). Equation (4) reduces to

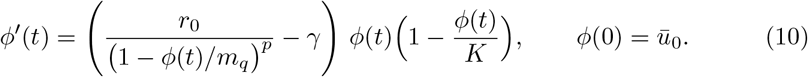

The singularity now occurs at *ϕ* = *m*_*q*_, and a standard Kawarada analysis [10, 22] yields the quenching-time asymptotics described below.

##### Monotonicity

Assume 0 *< ū*_0_ *<* min *K, m*_*q*_ and *r*_0_ *> γ*, so that the net growth is initially positive. The reduced ODE reads

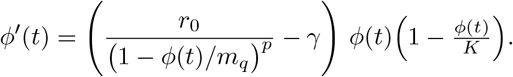

For any *ϕ* ∈ (0, min*{K, m*_*q*_*}*) the factor *ϕ*(1 − *ϕ/K*) is strictly positive, and the coefficient

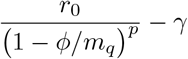

is minimized at *ϕ* = 0, where it equals *r*_0_ − *γ >* 0, and increases as *ϕ* ↑ *m*_*q*_ because the singular factor (1 − *ϕ/m*_*q*_)^−*p*^ is increasing. Hence

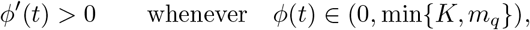

so *ϕ* is strictly increasing as long as it stays in this interval. In particular, *ϕ* increases from *ϕ*(0) = *ū*_0_ up to *m*_*q*_.

##### Finite-time hitting of m_q_ (quenching)

To prove finite-time quenching we follow an an alogous approach as in [10–13]. Indeed, fix any *ϕ*_*_ ∈ (0, *m*_*q*_), and write *g*(*ϕ*) := *ϕ*(1 − *ϕ/K*). On the compact interval [*ϕ*_*_, *m*_*q*_],

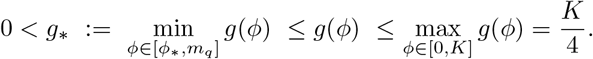

From (10),

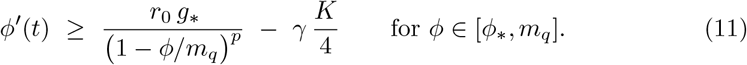

Choose *η* ∈ (0, 1) so close to 1 that for all *ϕ* ∈ [*m*_*q*_(1 − *η*), *m*_*q*_) we have

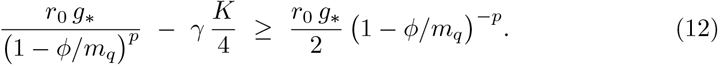

(For instance, it suffices to pick *η* so that 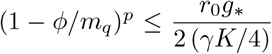 on that interval; this is always possible since the LHS can be made arbitrarily small when *ϕ* ↑ *m*_*q*_.) Let *t*_1_ be the first time with *ϕ*(*t*_1_) = *m*_*q*_(1 − *η*). Standard ODE comparison (or simply *ϕ*^*′*^(*t*) *>* 0) shows *t*_1_ *<* ∞.

For *t* ≥ *t*_1_ we combine (11)–(12) to obtain

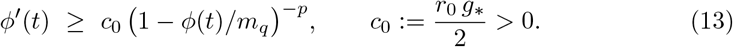

Let *ψ*(*t*) := *m*_*q*_ − *ϕ*(*t*) ∈ (0, *m*_*q*_*η*]. Then *ψ*^*′*^(*t*) = −*ϕ*^*′*^(*t*) ≤ −*c*_0_ (*ψ*(*t*)*/m*_*q*_)^−*p*^ = −*c ψ*(*t*)^−*p*^ with 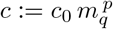. Hence

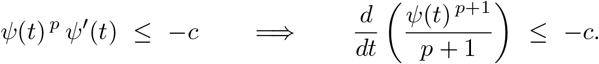

Integrating on [*t*_1_, *t*) gives

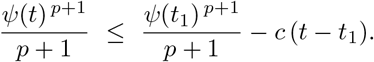

Therefore *ψ*(*t*) ↓ 0 in finite time, and the *quenching time* satisfies the explicit upper bound

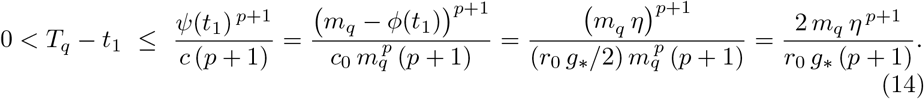

Thus *ϕ*(*t*) ↗ *m*_*q*_ as *t* ↑ *T*_*q*_ *<* ∞, while

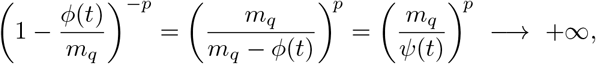

and, by (9), the reaction term *F* (*ϕ*; *w*) diverges. This is *finite-time quenching of Kawarada type* for the spatially homogeneous solution.

##### Explicit asymptotics

Keeping only the dominant term in (10) as *ϕ* ↑ *m*_*q*_,

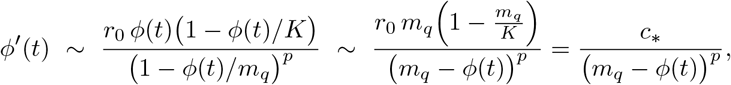

with *c*_*_ := *r*_0_ *m*_*q*_ (1 − *m*_*q*_*/K*) *>* 0 (since *m*_*q*_ ≤ *K* by the choice of initial data and parameters). Then

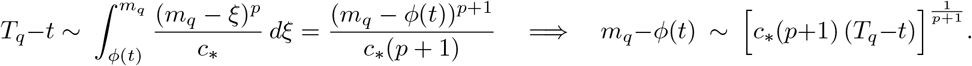

Consequently

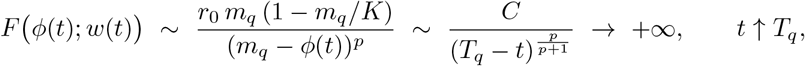

confirming the quenching rate.

#### Extension to nonhomogeneous initial data

The homogeneous reduction (10) provides a certificate that the full nonlocal Kawarada PDE admits quenching solutions under Neumann boundary conditions. This extends to a class of nonhomogeneous initial data by a comparison argument. Let *ū*_0_ ∈ (0, min {*K, m*_*q*_}) be such that the solution *ϕ* of (10) quenches at time *T*_*q*_, and consider the corresponding spatially homogeneous solution *v*(*x, t*) = *ϕ*(*t*). If the initial profile 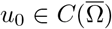 satisfies

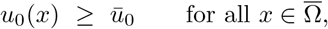

then *v* is a classical subsolution of the PDE with Neumann boundary conditions, while the true solution *u* with initial data *u*_0_ is a supersolution. The nonlocal reaction term is monotone in *u* thanks to the positivity of the kernel and of the logistic factor, so the parabolic comparison principle applies and yields

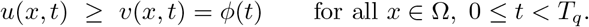

In particular,

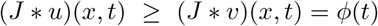

for all *x* ∈ Ω and *t < T*_*q*_. Since *ϕ*(*t*) ↗ *m*_*q*_ as *t* ↑ *T*_*q*_, the nonlocal quantity (*J* * *u*)(*x, t*) is driven towards the same critical threshold in the sense that

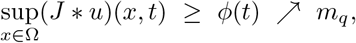

so the nonlocal factor 1 − (*J* * *u*)(*x, t*)*/m*_*q*_ becomes arbitrarily small near *t* = *T*_*q*_.

Because *u*(·, *t*) remains bounded and strictly positive on any compact subset of Ω for *t < T*_*q*_ (by the strong maximum principle and the fact that *u*_0_ ≥ *ū*_0_ *>* 0), the logistic prefactor *u*(*x, t*) 1 − *u*(*x, t*)*/K* is bounded away from zero on a subset of nonzero measure. On this subset, as *t* ↑ *T*_*q*_ we have

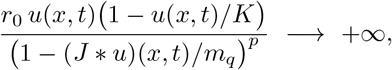

so the reaction term diverges and the right-hand side of the PDE

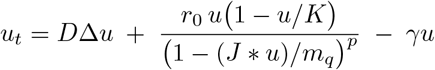

blows up in finite time. The diffusion term *D*Δ*u* and the linear decay term −*γu* remain bounded as *t* ↑ *T*_*q*_, whereas the singular nonlocal term dominates. Consequently,

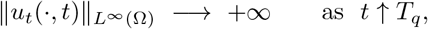

and the solution *u* quenches no later than *T*_*q*_ in the Kawarada sense: the solution itself remains bounded, but its time derivative and the reaction term blow up in finite time. Thus any initial condition that dominates a quenching homogeneous profile leads to finite-time quenching, with blow-up of *u*_*t*_, for the full nonlocal Neumann problem.

#### Biological interpretation

From a biological perspective, the spatially homogeneous reduction describes a tumour whose cell density is uniform in space and whose effective proliferation rate is still governed by a nonlocal measure of the tumour burden. In the homogeneous setting this nonlocal signal reduces to the scalar *ϕ*(*t*), which can be interpreted as the average cell density (or, up to a factor, the total tumour mass). The differential inequality (13) shows not only that *ϕ* is strictly increasing, but also that its time derivative diverges as *ϕ*(*t*) ↗ *m*_*q*_: along any quenching solution one has *ϕ*(*t*) → *m*_*q*_ and (1−*ϕ*(*t*)*/m*_*q*_)^−*p*^ → ∞ as *t* ↑ *T*_*q*_, hence *ϕ*^*′*^(*t*) → ∞. Thus, as the average density approaches the critical level *m*_*q*_, the instantaneous growth rate (or “velocity”) of the tumour explodes in finite time. This is precisely the continuum manifestation of an *explosive* growth regime: the tumour mass increases ever more rapidly as it fills the available tissue and approaches its effective capacity. This behaviour is consistent with the superlinear scaling patterns and accelerating growth dynamics reported for human cancers in [1], where empirical growth exponents *β >* 1 indicate that larger tumours proliferate disproportionately faster than smaller ones.

The comparison argument of the previous subsection shows that this picture is not restricted to perfectly homogeneous initial data. If the initial profile *u*_0_ dominates a quenching homogeneous state, then the corresponding solution *u*(*x, t*) of the full nonlocal Neumann problem remains bounded and strictly positive for *t < T*_*q*_, but we still have

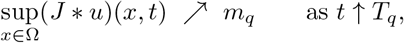

and the singular factor 1 − (*J* * *u*) (*x, t*)*/m* ^−*p*^ diverges on a set of nonzero measure. In particular, on regions where *u*(*x, t*) does not degenerate, the nonlocal proliferation term

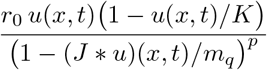

blows up, and the diffusion and linear decay terms remain bounded. As a consequence, the time derivative *u*_*t*_ itself blows up in finite time, even for nonhomogeneous profiles: the tumour density stays below saturation, but its rate of change becomes unbounded. In this sense, both the homogeneous and the nonhomogeneous cases exhibit the same qualitative scenario of explosive growth driven by a nonlocal feedback.

The quenching rates

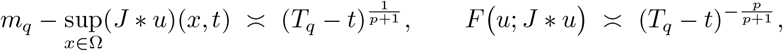

express how the system approaches this explosive regime, with the nonlocal environment *J * u* controlling the proximity to the critical threshold and the reaction term providing a measure of the instantaneous growth speed. Although our model is more singular than the empirical power laws in [1], it illustrates rigorously how nonlocal, size-dependent feedback can drive the system toward a finite-time singularity in growth speed, in qualitative agreement with the accelerated growth patterns seen in PET and MRI data.

#### Link between *p* and *β*

The universal scaling law reported in [1] describes an empirical, cross–sectional allometric relation between total lesion activity (TLA) and metabolic tumour volume (MTV) of the form

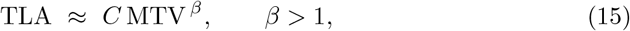

where *β* characterizes a population-level size–activity superlinearity measured at a fixed observation time. In contrast, the exponent *p >* 0 appearing in the Kawaradatype singular reaction term

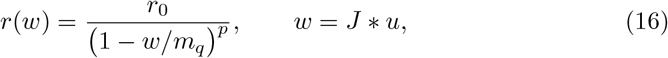

governs a temporal acceleration mechanism: as the nonlocal field *w* approaches the critical threshold *m*_*q*_, the reaction rate diverges like (*m*_*q*_ − *w*)^−*p*^ (up to a multiplicative constant), potentially driving finite-time explosive dynamics.

It is important to emphasize at the outset that there is, in general, no model-independent identity of the form *p* = *f* (*β*). The exponent *β* describes how activity scales with tumour volume across a cohort at a given observation level, whereas *p* controls how rapidly the mechanistic reaction rate diverges in time as the system approaches a singular threshold. A quantitative link between the two exponents therefore requires an additional modelling closure connecting the distance-to-singularity in (16) to the imaging observables (MTV,TLA).

To clarify the temporal role of *p*, consider an accelerating regime in which the system approaches the singular time *T*_*q*_. Let

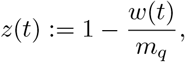

so that *z*(*t*) ↓ 0 corresponds to *w*(*t*) ↑ *m*_*q*_. Retaining only the dominant singular contribution in the reaction term and treating the remaining multiplicative factors as slowly varying in this regime leads formally to a reduced balance of the type

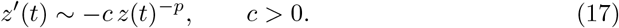

This classical singular ordinary differential equation yields the asymptotic scaling

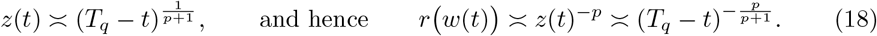

Accordingly, the exponent *p* determines the temporal acceleration rate 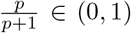 governing the divergence of the mechanistic reaction term as *t* ↑ *T*_*q*_.

To relate this temporal singularity mechanism to the cross–sectional allometry (15), one may introduce a minimal observational closure in an accelerating regime. A parsimonious ansatz is

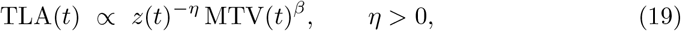

which expresses two effects simultaneously: first, that activity increases with tumour burden according to the empirically observed exponent *β*, and second, that activity is further amplified as the system approaches the singular threshold *z*(*t*) ↓ 0. Here *β* is used as a phenomenological proxy for size-dependence along a typical trajectory, rather than being derived from the PDE. Combining (19) with (18) yields

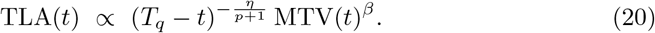

Under this closure, *p* controls the temporal amplification factor (*T*_*q*_ − *t*)^−*η/*(*p*+1)^, whereas *β* continues to describe an instantaneous size–activity scaling relation.

A stronger, yet more restrictive, identification may be obtained by assuming that the measured activity is proportional to the effective singular rate acting on active volume, namely

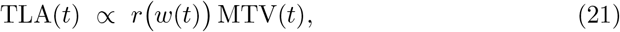

which corresponds to interpreting the mean metabolic intensity per unit volume as being proportional to the reaction rate *r*(*w*). Substituting (18) into (21) gives

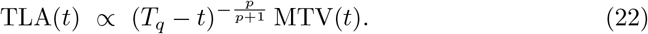

Combining (22) with the empirical allometry (15) yields the constraint

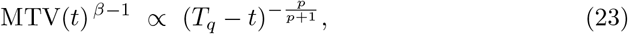

which shows that a direct relation between *p* and *β* still requires an additional description of the time evolution of MTV(*t*) near *T*_*q*_. For instance, if MTV(*t*) ≍ (*T*_*q*_ − *t*)^−*ζ*^ for some *ζ >* 0, then

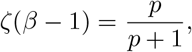

leading to a conditional identification of *p* in terms of *β* and the mass-growth exponent *ζ*. Absent such a dynamical closure, no standalone identity *p* = *f* (*β*) can be deduced.

In summary, the exponents *p* and *β* both quantify superlinearity, yet they act on fundamentally different aspects of the system. The parameter *p* governs temporal acceleration toward a mechanistic singularity in the nonlocal field *w* = *J* ∗ *u*, whereas *β* characterizes an instantaneous population-level scaling between (TLA,MTV). A quantitative bridge between them becomes possible only after specifying an explicit observation model linking the singular dynamics of the PDE to imaging-derived quantities.

### 3.3 Stability and travelling waves

In this section we analyse the dynamics of small perturbations around steady states of (4). The resulting linear stability analysis yields spectral conditions that determine whether such perturbations decay or grow, and thus whether the corresponding tumour configurations are asymptotically stable, unstable, or prone to pattern-forming instabilities within the nonlocal model. From a biological standpoint, this characterises whether small spatial or temporal variations in cell density are damped out—leading to robust, homogeneous tumour growth—or instead amplified, potentially giving rise to heterogeneous, aggressively expanding tumour structures. In the one-dimensional setting we complement this with a travelling-wave analysis, where coherent invasion fronts connect different steady states and provide an effective description of directed tumour spread through tissue, with the wave speed and profile encoding how fast and in what manner the cancer infiltrates the surrounding environment under the influence of nonlocal proliferation.

#### Frécet linearization around a general steady state

Let *u*_*s*_(*x*) be a (sufficiently smooth) steady state of (4), i.e. a solution of

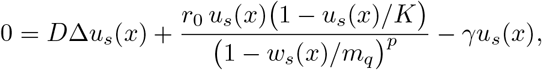

where

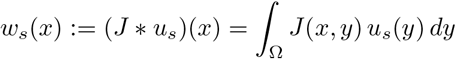

denotes the associated nonlocal density.

Recall that from (8) we can write the PDE in the compact form

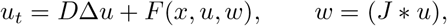

where

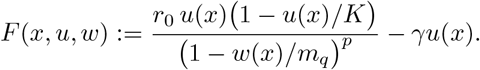

To linearize about *u*_*s*_ we set

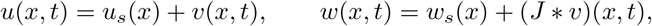

and expand *F* to first order in *v*. Using the chain rule we obtain

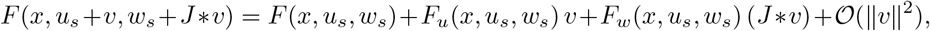

where subscripts denote (Frécet) partial derivatives with respect to the second and third aguments of *F*.

Since *u*_*s*_ is a steady state, *F* (*x, u*_*s*_, *w*_*s*_) = − *D*Δ*u*_*s*_(*x*) and cancels with the diffusion term in the PDE. Keeping only linear terms in *v* yields

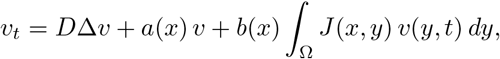

with

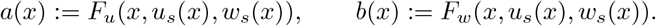

We now compute *a*(*x*) and *b*(*x*) explicitly. Differentiating *F* with respect to *u* gives

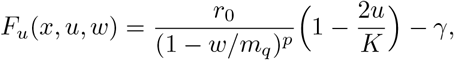

and differentiating with respect to *w* yields

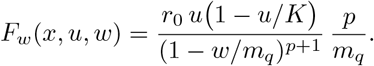

Evaluating at (*u*_*s*_(*x*), *w*_*s*_(*x*)) we obtain

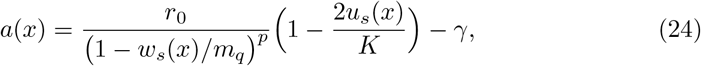

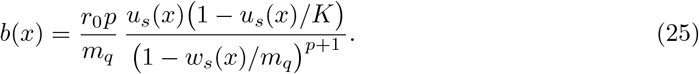

This defines a nonlocal linearized operator

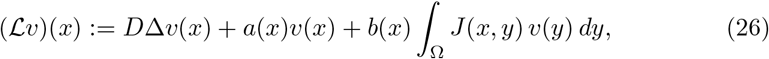

with Neumann domain

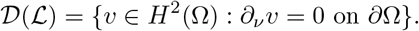

The linear stability of *u*_*s*_ is determined by the spectral bound

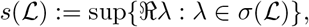

where *σ*(ℒ) denotes the spectrum of ℒ and ℜ*λ* stands for the real part of the eigenvalue *λ*. If *s*(ℒ) *<* 0 then small perturbations decay exponentially and *u*_*s*_ is linearly asymptotically stable; if *s*(ℒ) *>* 0 then there exist perturbations that grow exponentially and *u*_*s*_ is linearly unstable.

#### Spatially homogeneous steady states

A particularly important class of steady states in the Neumann setting is the spatially homogeneous ones. Assume *u*_*s*_(*x*) ≡ *ū* is constant. Using the normalization (K2), for each *x* ∈ Ω we have

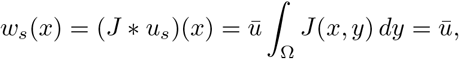

so *w*_*s*_ is also constant. In this case, (24)–(25) simplify to constants *a*_*N*_, *b*_*N*_ :

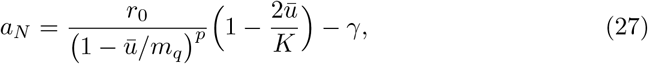

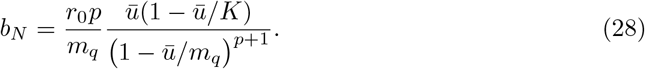

The linearized equation becomes

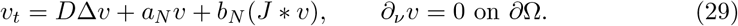

Let {−*µ*_*k*_}_*k*≥0_ denote the Neumann eigenvalues of −Δ on Ω, with corresponding orthonormal eigenfunctions {*ϕ*_*k*_}_*k*≥0_:

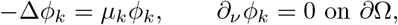

where 0 = *µ*_0_ ≤ *µ*_1_ ≤ *µ*_2_ ≤ · · ·, *µ*_*k*_ → ∞. Any perturbation can be expanded as

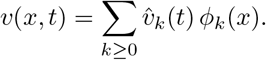

The action of the convolution operator can be written as

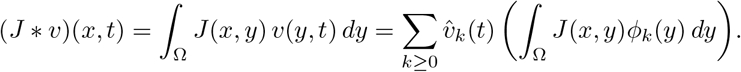

In general, the nonlocal term couples the modal amplitudes 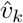. To obtain a more transparent spectral description, we now restrict to kernels *J* that are of convolution type with respect to an underlying geometry for which the Laplacian eigenfunctions form a Fourier-like basis. Concretely, we assume that there exists a family of eigenfunctions {*ϕ*_*k k*≥0_} of the Neumann Laplacian (for example cosine modes on an interval, or trigonometric modes on a periodic box) such that *J* depends only on the relative position of its arguments in this geometry, e.g. *J*(*x, y*) = 𝒥 (*x* − *y*) or *J*(*x, y*) = 𝒥 (dist(*x, y*)), see Fig. 2

**Figure 2.**
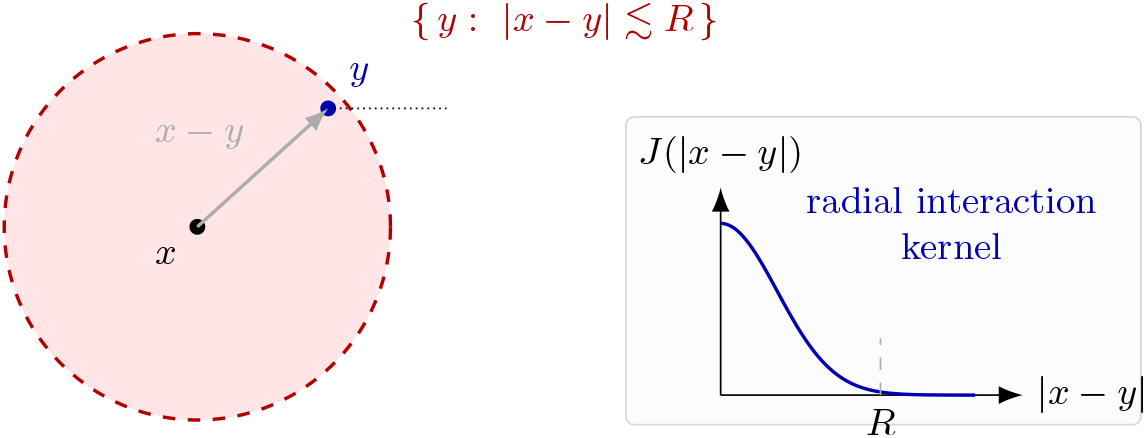
Geometry of a radial nonlocal convolution kernel. Points *x* and *y* interact within a neighbourhood of radius *R*, and the kernel depends only on the distance |*x* − *y*|. The inset shows a typical radial profile *J* (|*x* − *y*|).

In that case the integral operator

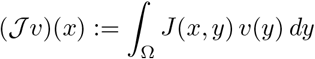

is diagonal in the basis {*ϕ*_*k k*≥0_}, i.e. each Laplacian eigenfunction is also an eigenfunction of the nonlocal operator. More precisely, we assume

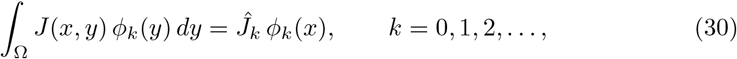

for some real numbers *Ĵ*_*k*_, which play the role of Fourier coefficients (or the spectral transform) of the kernel *J* in the chosen geometry. Under this assumption the nonlocal term acts modewise, and the linearized dynamics can be analysed separately for each mode *k*. Substituting the modal expansion into (29) and using (30) gives

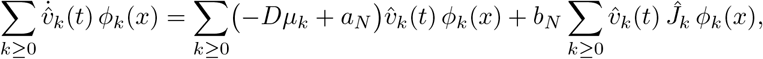

so that each mode satisfies the decoupled ODE

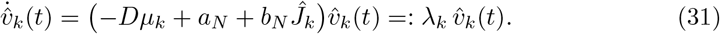

Thus the eigenvalues of the linearized operator restricted to the homogeneous steady state are

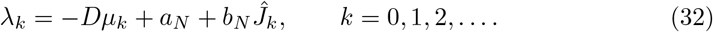

The spatially homogeneous steady state *u*_*s*_ ≡ *ū* is linearly asymptotically stable if

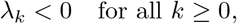

and unstable if there exists at least one mode *k* with *λ*_*k*_ *>* 0. Notice that *µ*_0_ = 0 and *ϕ*_0_ is constant; the corresponding eigenvalue *λ*_0_ controls the stability of spatially homogeneous perturbations, while the higher modes (*k* ≥ 1) describe pattern-forming instabilities. The nonlocal term *b*_*N*_ *Ĵ*_*k*_ couples the kernel and the singularity exponent *p* to these stability properties: if *p* is large and *Ĵ*_*k*_ *>* 0 for some *k* ≥ 1, the term *b*_*N*_ *Ĵ*_*k*_ can compensate the stabilizing diffusion −*Dµ*_*k*_ and destabilize the homogeneous state.

#### Travelling wave analysis in one dimension

We now turn to travelling wave solutions in one spatial dimension, *x* ∈ ℝ, for which the domain is unbounded and Neumann boundary conditions correspond to asymptotically flat profiles. In this setting we assume that the nonlocal kernel is translation-invariant, i.e.

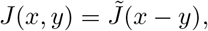

for some one-dimensional profile 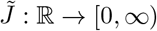 normalized by 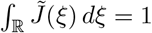, cf. Fig. 2 . Thus 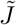 is the one-dimensional convolution kernel associated with the two-point kernel *J* used in the bounded-domain analysis.

We then look for travelling-wave solutions of the form

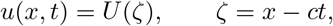

where *c* ∈ ℝ is the wave speed (typically *c >* 0 for a right-moving front).

##### Derivation of the travelling-wave equation

With the travelling-wave ansatz *u*(*x, t*) = *U* (*ζ*) we compute

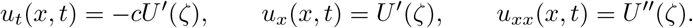

The nonlocal term becomes

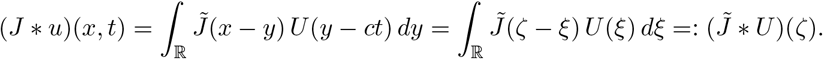

Substituting into (4) (with Ω = ℝ and kernel 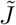) yields the travelling-wave ODE

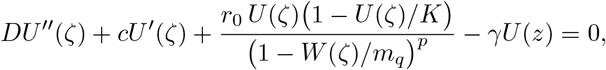

where 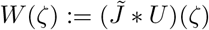 stands for the nonlocal density along the wave profile.

##### Asymptotic states and boundary conditions

We are interested in fronts connecting two (possibly distinct) homogeneous steady states *U*_−_, *U*_+_:

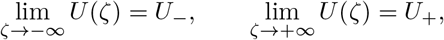

with *U*_*±*_ constant and satisfying the steady-state equation

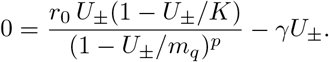

Because 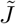 is normalized, *W* (*ζ*) → *U*_*±*_ as *ζ* → *±*∞. The Neumann (no-flux) condition on a large but finite interval [−*L, L*] can be approximated by

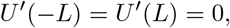

which corresponds, in the whole-line limit *L* → ∞, to asymptotically flat profiles

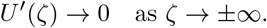

##### Linearization at asymptotic states and dispersion relation

To understand the local structure of a travelling wave near its asymptotic states, we linearize the comoving equation at a constant equilibrium *U*_∗_ (either *U*_−_ or *U*_+_). Recall that in the travelling frame *ζ* = *x* − *ct* the profile *U* satisfies

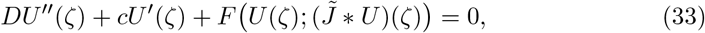

recalling that

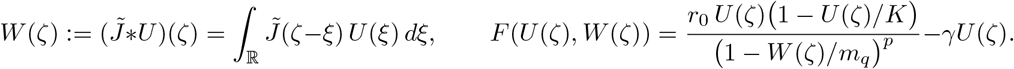

A constant equilibrium *U*_∗_ satisfies *F* (*U*_∗_, *U*_∗_) = 0; the asymptotic states *U*_*±*_ are examples of such equilibria.

We perturb *U*_∗_ by writing

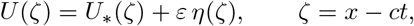

and define

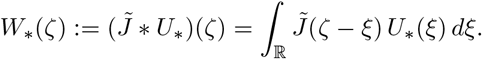

Since *U*_∗_ is constant and 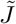 is normalized, we have *W*_∗_(*z*) *U*_∗_. For the perturbation we obtain

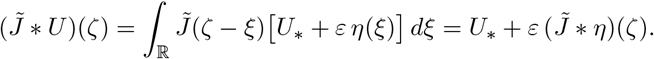

Substituting *U* = *U*_∗_ + *εη* and 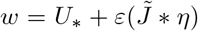 into (33) and expanding *F* to first order in *ε* yields

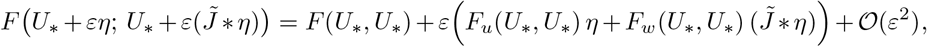

where *F*_*u*_ and *F*_*w*_ denote Frécet partial derivatives with respect to *u* and *w* respectively, evaluated at (*U*_∗_, *U*_∗_). Since *F* (*U*_∗_, *U*_∗_) = 0, the 𝒪 (1) term vanishes. Collecting the 𝒪 (*ε*) terms and dropping higher-order contributions gives the linearized equation

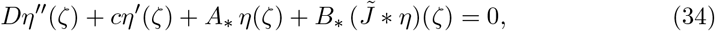

where

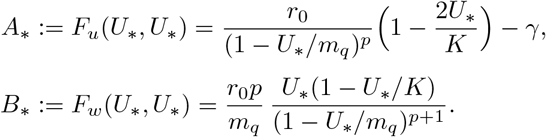

These constants coincide with *a*_*N*_, *b*_*N*_ in (27)–(28) evaluated at *ū* = *U*_∗_ and describe the local linear response of the nonlocal reaction term about the equilibrium *U*_∗_.

To extract the dispersion relation, we look for exponential solutions of the form

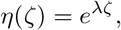

so that

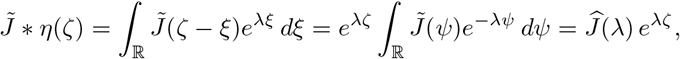

with

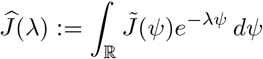

the (two-sided) Laplace transform of 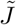. Substituting into the linearized equation (34) yields

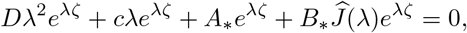

and, after cancelling *e*^*λζ*^*≠* 0, the dispersion relation

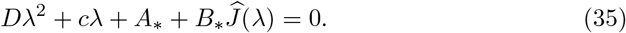

For a given asymptotic state *U*_∗_ and kernel 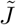, admissible wave speeds *c* must be such that (35) has roots *λ* with the correct sign of ℜ*λ* to enforce the desired exponential decay as *z* → *±*∞. Typically, for a wave connecting *U*_−_ to *U*_+_, one requires

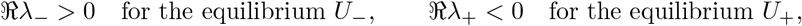

so that perturbations decay in the respective far fields. The singular exponent *p* enters via *A*_∗_ and *B*_∗_ and thereby influences both the admissible wave speeds and the sharpness of the travelling front, linking the strength of the nonlocal singular feedback to the invasion dynamics described by the wave. From the viewpoint of tumour growth, increasing *p* corresponds to stronger acceleration of proliferation as the local nonlocal burden approaches the critical threshold, which in turn can lead to faster, steeper invasion fronts and more aggressive tissue infiltration in the model. A more detailed travelling-wave analysis will be presented in a forthcoming paper.

## 4 Bayesian inference framework for the nonlocal Kawarada-type model

The nonlocal reaction–diffusion model (4) provides a mechanistic description of tumour cell density evolution in which proliferation is modulated by a spatially aggregated nonlocal field *w*(*x, t*) = (*J* ∗ *u*)(*x, t*) and exhibits Kawarada-type acceleration as *w* approaches a critical threshold *m*_*q*_. From an inferential perspective, the principal goal is to quantify the uncertainty in the parameters governing diffusion, logistic saturation, decay, and singular amplification, while ensuring that the resulting inference remains consistent with the limited observational resolution available in clinical imaging datasets. In the present study, the primary imaging observables are patient-level summaries such as metabolic tumour volume (MTV) and total lesion activity (TLA), which are cross–sectional in nature and do not directly resolve the spatial field *u*(*x, t*) or its temporal evolution. A Bayesian framework is therefore attractive because it allows one to combine mechanistic structure with explicit observation models and scientifically motivated priors, and it makes transparent which components of the parameter vector are data-informed and which remain prior-dominated under the available measurements. In particular, any literature-based values reported for cross– sectional exponents (e.g. *β* = 5*/*4 in [1]) are treated here as *benchmarks for comparison* rather than hard constraints: the scaling exponent is inferred from the cohort and its posterior uncertainty is reported.

To emphasise this point, it is convenient to distinguish between two related but conceptually different inferential targets. The first target is an observation-level description of the cross–sectional relationship between TLA and MTV, as captured by the universal scaling law (15). The second target is a mechanistic interpretation of the observed superlinearity in terms of the singular exponent *p* and other PDE parameters in (4). The Bayesian framework developed below is designed to accommodate both targets in a coherent manner by separating the *forward model* (mechanistic dynamics) from the *observation model* (mapping of dynamics to imaging-derived quantities) and by explicitly encoding a closure that bridges the PDE parameters to the cross–sectional scaling exponent.

### 4.1 Model specification and parameterization

We consider the nonlocal Kawarada-type reaction–diffusion model (4) and collect the mechanistic parameters into the vector

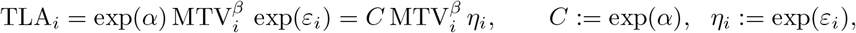

where *ϑ*_*J*_ denotes the parameter(s) governing the kernel *J*. In applications where *J* is taken to be Gaussian, one may write

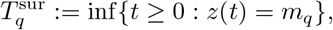

so that *ϑ*_*J*_ = *ℓ >* 0 represents the interaction length-scale and the normalization ensures condition (K2). The inference problem consists of learning *θ* from patient-level observations, while accounting for measurement noise and structural mismatch between the continuum model and clinical data.

### 4.2 Observation model for cross–sectional imaging data

Let (MTV_*i*_, TLA_*i*_) denote the paired PET-derived summaries for patient *i* = 1, …, *n* recorded at a clinically defined observation visit. Since the underlying nonlocal FKPP-type model (4) evolves a spatial field, an observation operator is, in principle, required to map the latent state to imaging summaries. A natural class of operators is obtained by integrating suitable functionals of the tumour field over a patient-specific domain Ω_*i*_. For instance, if *u*_*i*_(·, *t*) denotes a normalized tumour cell-density field, then a canonical volume proxy is the total mass

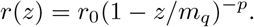

and an activity proxy may be represented as an intensity-weighted integral

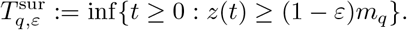

for a suitable metabolic activity density *a*(·,·) and nonlocal field *w*_*i*_. Although such operators are scientifically meaningful, they are not identifiable from a single cross– sectional pair (MTV_*i*_, TLA_*i*_) without longitudinal imaging or spatially resolved PET information. To remain aligned with the empirical strategy of [1] while retaining a mechanistic connection to the nonlocal FKPP parameters, we therefore adopt a reduced observation model in which the PDE state is summarized by a low-dimensional surrogate and the cross–sectional scaling relation is modelled directly on the log scale.

We model the cross–sectional MTV–TLA allometry on the log scale via

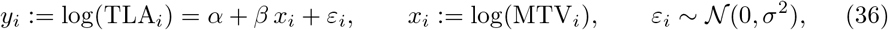

where *α* is an intercept and *β* is the scaling exponent. Equivalently, exponentiating (36) yields the multiplicative power-law representation

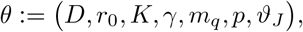

so that *C* is the allometric prefactor and *η*_*i*_ captures multiplicative noise on the original scale while *β* is the scaling exponent in the universal scaling law (15), and *σ*^2^ collects measurement noise and model discrepancy on the log scale. We estimate *β* from the cohort; when plotting or discussing results we may also indicate the reference value *β* = 5*/*4 from [1] solely as a visual/interpretive benchmark, not as an imposed constraint. In our Bayesian implementation, the log–log regression is used to estimate and summarize the cross–sectional scaling behaviour, while the mechanistic model is simultaneously constrained by additional observation equations that link MTV and TLA to a latent progression state.

### 4.3 Reduced mechanistic surrogate and latent stage times

To bridge the nonlocal FKPP mechanism to cross–sectional summaries without patient-specific spatial domains, we introduce a reduced mean-field surrogate *z*(*t*) ∈ (0, 1) that summarizes the tumour progression state. The surrogate is defined implicitly as the solution of an ODE with parameters inherited from the mechanistic model, and it is evaluated on a fixed temporal grid *t* ∈ [0, *T*_max_].

In the mechanistic model (4), the quenching time *T*_*q*_, defined in Definition 2, is the first time at which the latent progression state reaches the singular threshold *m*_*q*_. In the present reduced surrogate formulation, the analogous quantity is

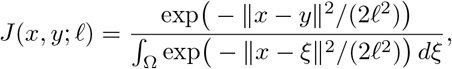

which corresponds to the onset of singular amplification in the growth factor

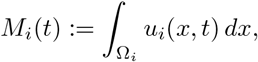

By contrast, *T*_max_ is a user-chosen finite time horizon used to define the numerical grid for solving the reduced dynamical system and to constrain the latent stage times via *t*_*i*_ ∈ (0, *T*_max_). In principle, one chooses *T*_max_ large enough to cover the pre-quenching regime of interest, for example with 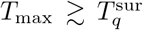 for typical parameter values, and the intended interpretation is that observations occur before quenching, that is, 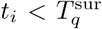. In the implementation, we regularize the singularity by replacing 1 −*z/m*_*q*_ with max{1 −*z/m*_*q*_, *ε*} for a small *ε >* 0, so that a practical quenching proxy is

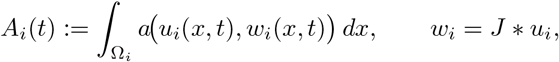

Each patient is associated with an unobserved stage time *t*_*i*_ ∈ (0, *T*_max_) representing the latent disease stage at which the PET measurement is realized. Given (*t*_*i*_), we define *z*_*i*_ := *z*(*t*_*i*_) and connect *z*_*i*_ to the observed MTV and TLA through multiplicative log-normal error models.

Recall that *m*_*q*_ ∈ (0, 1] denote the nonlocal threshold parameter and *p >* 0 the singularity exponent, we define the singular amplification factor by

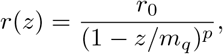

with *r*_0_ *>* 0. The progression dynamics in *z* are governed by a logistic-type modulation with parameters *γ >* 0 and *K* ∈ (0, 1], yielding an ODE of the form

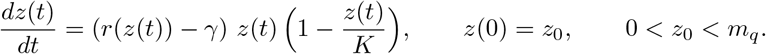

This reduced surrogate should be interpreted as a cohort-level summary of the nonlocal FKPP dynamics rather than a full spatial reconstruction; it is nevertheless sufficient to provide an explicit forward map from mechanistic parameters to latent progression states used in the observation model.

Conditional on *z*_*i*_ = *z*(*t*_*i*_), we model the imaging summaries by

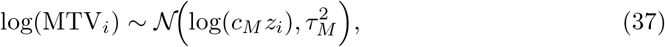

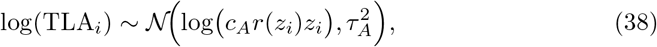

where *c*_*M*_ *>* 0 and *c*_*A*_ *>* 0 are cohort-level scaling constants and *τ*_*M*_, *τ*_*A*_ *>* 0 capture multiplicative measurement noise and residual model discrepancy for MTV and TLA, respectively. Together, (37)–(38) provide a mechanistic link between the non-local FKPP-type growth mechanism and the cross–sectional PET-derived summaries through the latent stage times (*t*_*i*_) and the surrogate trajectory *z*(·).

### 4.4 Bridge from mechanistic parameters to the scaling exponent

The singularity exponent *p* governs the strength of the acceleration mechanism through the factor (1 − *z/m*_*q*_)^−*p*^, whereas the scaling exponent *β* in (36) is an effective cross– sectional quantity. To allow the scaling law to inform the mechanistic calibration without requiring a model-independent identity, we introduce a low-dimensional closure relating *β* to (*p, ρ*), where *ρ* ∈ (0, 1) represents an effective cohort-level proximity to the threshold regime. In the implementation, the scaling exponent is represented as a monotone map

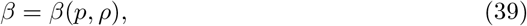

so that the regression (36) may be written as

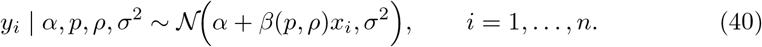

Equation (39) is interpreted as an explicit phenomenological bridge that captures the effect of singular acceleration on effective superlinearity while allowing uncertainty in the proximity parameter *ρ*.

### 4.5 Prior distributions and constraints

The prior specification reflects both the physical constraints of the reduced mechanistic system and the fact that several components are only indirectly informed under cross-sectional observation. In particular, the priors are chosen to respect the natural support of the parameters while remaining weakly informative, so that posterior learning is driven primarily by the observed MTV–TLA pairs rather than by restrictive prior assumptions.

The singularity exponent *p >* 0 is assigned a Gamma prior,

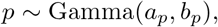

while the proximity parameter *ρ* ∈ (0, 1) is assigned a Beta prior,

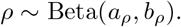

These choices are natural because the Gamma family is supported on the positive half-line and therefore matches the constraint *p >* 0, whereas the Beta family is supported on (0, 1) and is therefore well suited to parameters such as *ρ* that represent normalized proximity or fraction-type quantities.

We place a Gaussian prior on the log–log intercept,

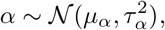

and half-normal priors on the scale parameters,

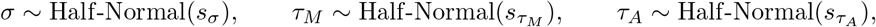

which is consistent with the computational implementation in which standard deviations are modelled directly rather than variances. The half-normal form is preferred here because it enforces positivity while mildly shrinking implausibly large noise levels without imposing an excessively heavy tail.

The remaining mechanistic and observation-scale parameters are likewise constrained by positivity or unit-interval bounds and are assigned weakly informative priors consistent with these constraints, for example

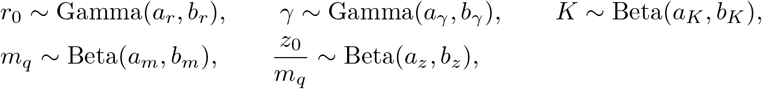

together with log-normal priors on *c*_*M*_ and *c*_*A*_, equivalently Gaussian priors on log *c*_*M*_ and log *c*_*A*_. The ratio prior on *z*_0_*/m*_*q*_ is used instead of assigning independent priors directly to *z*_0_ and *m*_*q*_ because it automatically enforces the biologically required constraint 0 *< z*_0_ *< m*_*q*_.

Finally, the latent stage times are constrained to (0, *T*_max_) and are assigned independent Beta priors on the normalized scale,

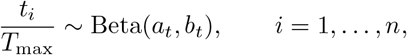

which provides a flexible way to encode weak prior information about typical observation stages while preserving the required support.

### 4.6 Posterior distribution and computation

Using the previously introduced log-transformed summaries (*x*_*i*_, *y*_*i*_), and letting *z*(·) denote the reduced surrogate trajectory determined by (*r*_0_, *γ, K, m*_*q*_, *p, z*_0_), the Bayesian model is defined by the scaling regression likelihood (40), the mechanistic observation likelihoods (37)–(38), and the priors in Section 4.5. Denoting **t** = (*t*_1_, …, *t*_*n*_)^⊤^, the joint posterior is proportional to

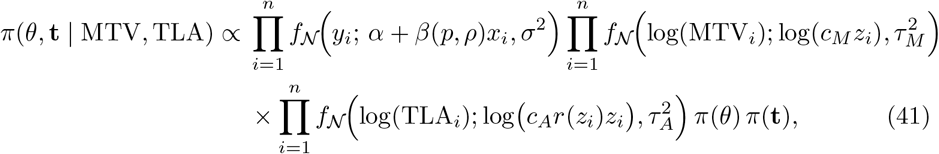

where *θ* collects all non-latent parameters, *z*(·; *θ*) is the surrogate solution, and *z*_*i*_ = *z*(*t*_*i*_; *θ*). Here *f*_𝒩_ (·; *µ, σ*^2^) denotes the Gaussian density with mean *µ* and variance *σ*^2^. Posterior computation is carried out using Metropolis–Hastings updates with transformed proposals that respect parameter constraints, including log transforms for positive parameters and logit transforms for unit-interval parameters. Since the likelihood evaluations depend on *z*(·; *θ*), each update of (*r*_0_, *γ, K, m*_*q*_, *p, z*_0_) entails recomputing the surrogate trajectory on the fixed grid, whereas updates of (*t*_*i*_) require only interpolation of the already-computed trajectory. Posterior summaries of interest include the marginal posteriors for the mechanistic parameters, the derived posterior for the scaling exponent *β*(*p, ρ*), and posterior predictive assessments of the log–log scaling relation and the joint distribution of (MTV, TLA).

## 5 Data and statistical analysis

The dataset consists of paired measurements of Metabolic Tumour Volume (MTV) and Total Lesion Activity (TLA) for a cohort of breast cancer patients. MTV quantifies the extent of metabolically active tumour tissue, whereas TLA augments this information by weighting the active volume by its metabolic intensity. Together, these two biomarkers provide complementary information on tumour burden and are widely used in oncological studies.

The dataset exhibits substantial variability across patients. MTV values range from approximately 3.5 to more than 200 cm^3^, while TLA spans more than two orders of magnitude, from roughly 7 to 1800 SUV-integrated units. The relationship between the two biomarkers is visualised in Figure 3, where a log–log hexbin plot reveals a clear positive association together with notable heterogeneity in metabolic burden. This structure motivates the use of a bivariate measurement model in the Bayesian likelihood, including an explicit correlation term to capture residual dependence not explained by the mechanistic PDE dynamics. Furthermore, the wide dynamic range and strictly positive nature of both biomarkers justify the choice of a lognormal observational noise model.

**Figure 3.**
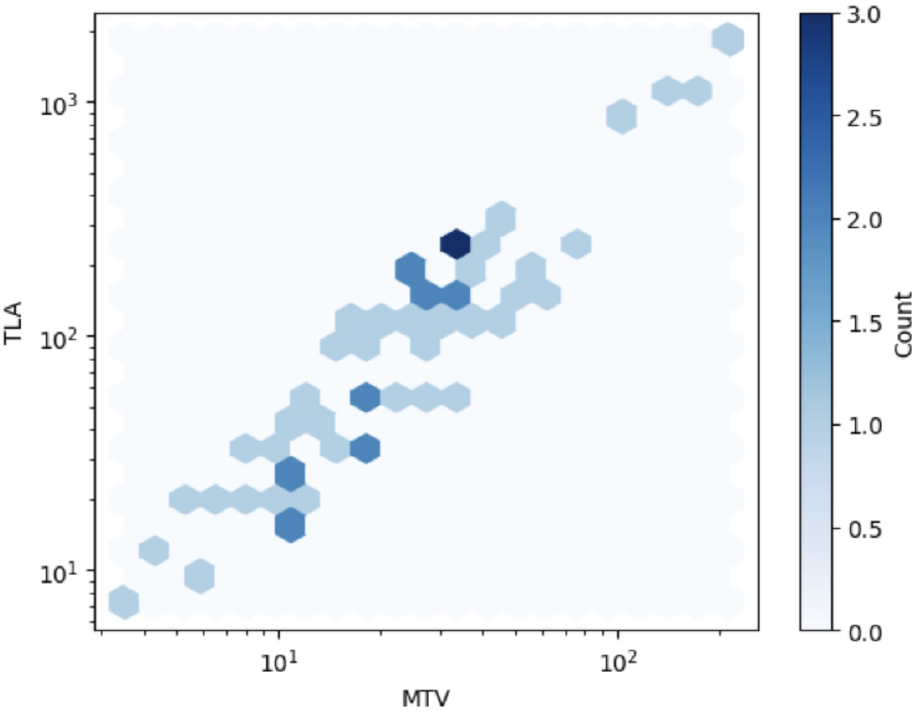
Log–log hexbin plot of MTV versus TLA for the breast cancer cohort, illustrating the strong positive association and wide dynamic range of metabolic burden.

The Bayesian calibration of the nonlocal FKPP-type tumour growth model to the paired MTV–TLA PET measurements provides a coherent and biologically plausible characterisation of metabolic tumour progression across the studied breast cancer cohort. The posterior means and 95% credible intervals summarise both the dominant scaling structure linking tumour activity to tumour burden and the residual heterogeneity that remains after accounting for this structure. A compact overview of posterior concentration across the principal parameters is given by the uncertainty summary in Figure 6, which indicates that some components are sharply informed by the data, whereas others remain comparatively diffuse, a pattern consistent with partial identifiability under cross-sectional observations.

The cross-sectional MTV–TLA relationship exhibits a clear log–log linear trend, and the posterior mean scaling fit tracks the central tendency of the data closely. The inferred scaling exponent is concentrated around *β* ≈ 1.30, with a 95% credible interval of approximately (1.20, 1.39), supporting superlinear scaling of total lesion activity with metabolic tumour volume across the observed range. The posterior distribution of *β* further indicates that the exponent is well identified in this cohort, as illustrated in Figure 4.

**Figure 4.**
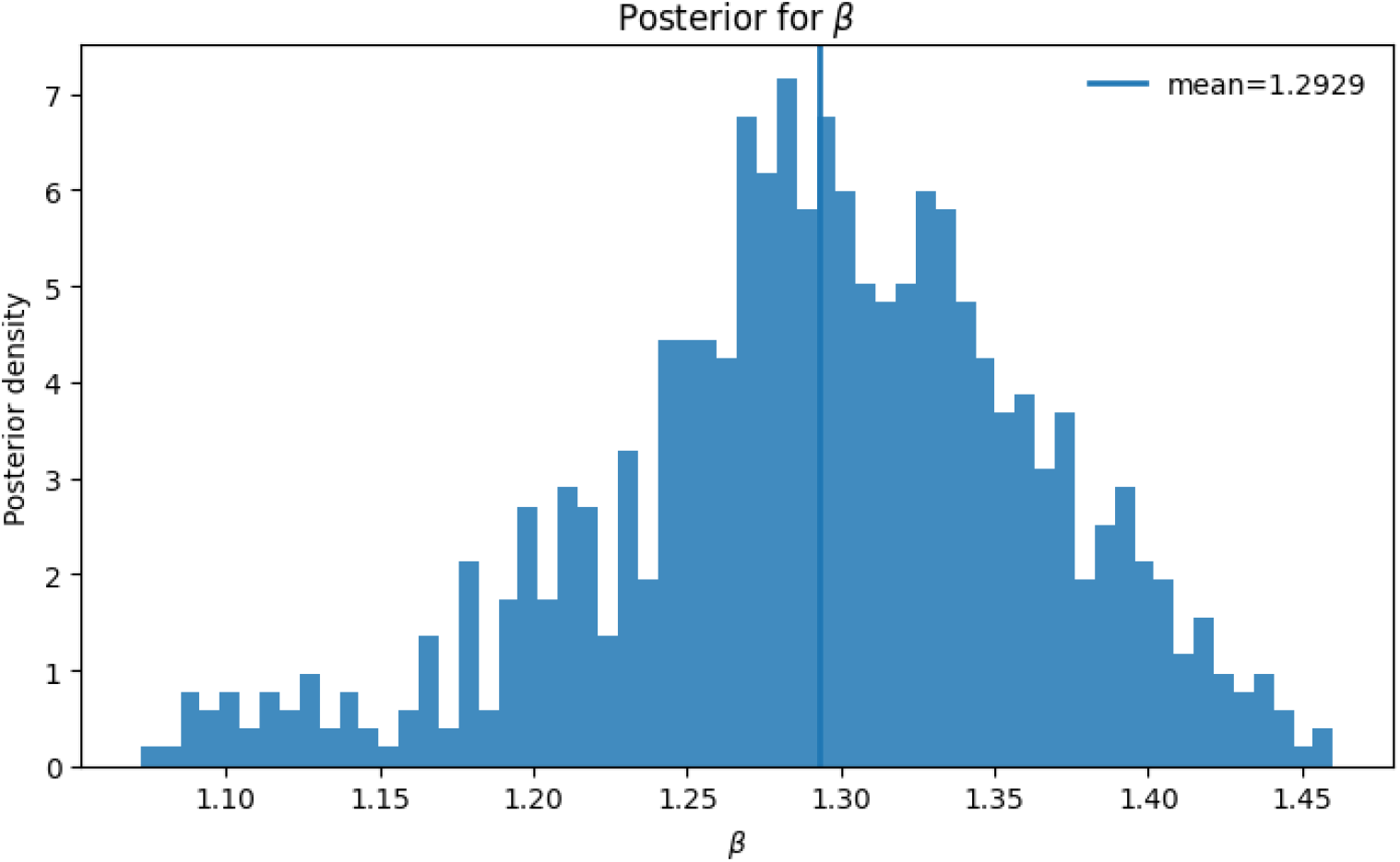
Cross-sectional MTV–TLA scaling: (a) log–log scatter with the posterior mean scaling fit; (b) posterior distribution of the scaling exponent *β*, with the posterior mean indicated by a vertical line.

The residual diagnostic provides an additional check of adequacy for the log–log regression component. The residuals plotted against log(MTV) are centred around zero without an obvious monotone pattern, which supports the constant-variance Gaussian error model on the log scale as a first-order approximation. The remaining dispersion is nonetheless substantial, consistent with biological and measurement heterogeneity not captured by MTV alone, and it motivates posterior predictive checks to verify that the assumed observation noise reproduces the empirical spread over the full range of tumour burden, cf. Figure 5.

**Figure 5.**
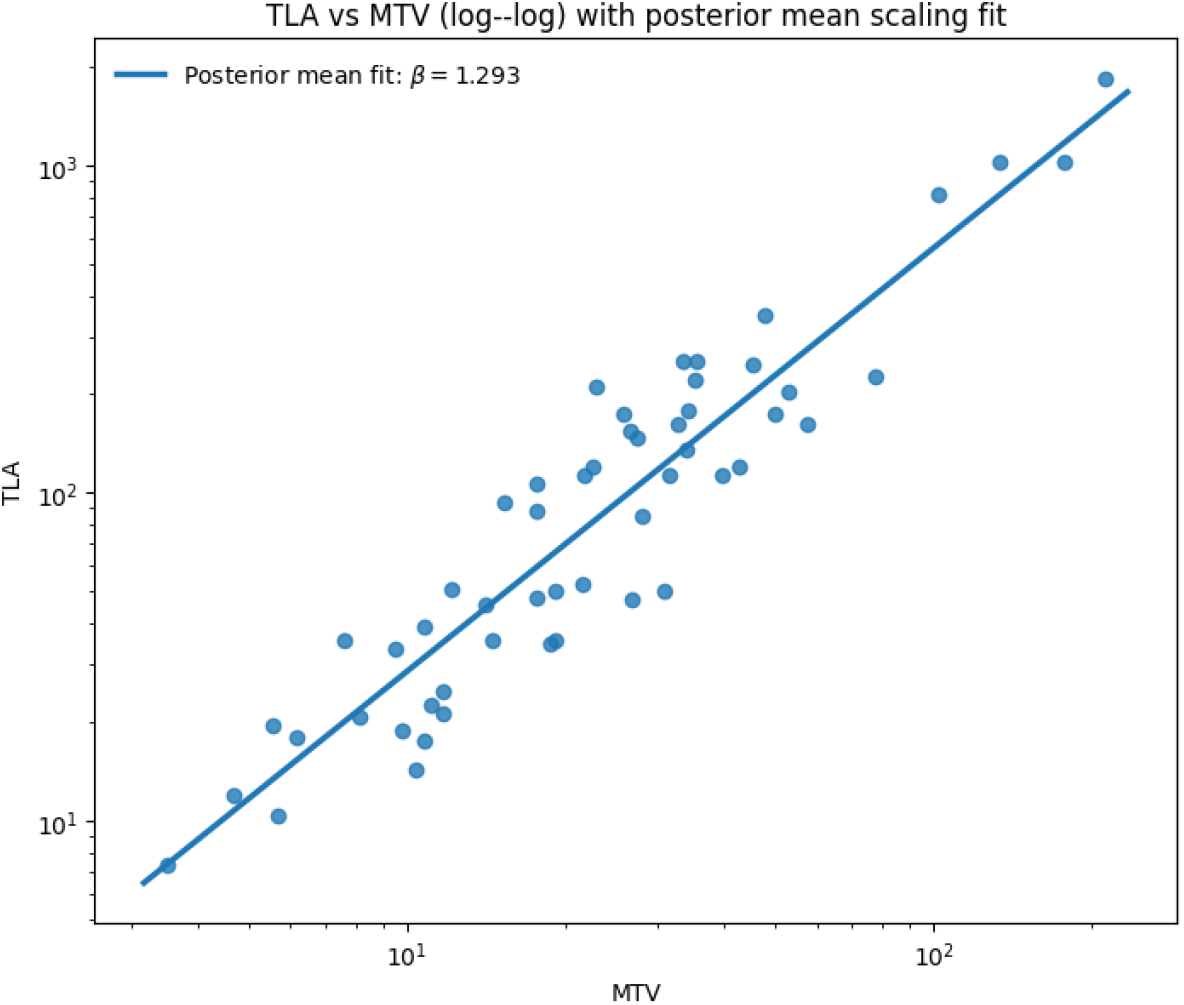
ℝesiduals in log(TLA) under the posterior mean log–log scaling fit, plotted against log(MTV).

Beyond the scaling fit, the posterior uncertainty is clearly heterogeneous across parameters, and this pattern is itself scientifically informative. Figure 6 shows that quantities tied most directly to the observation layer and the effective MTV–TLA scaling, in particular *β* and *σ*, are comparatively well constrained by the data, whereas several mechanistic parameters remain substantially more diffuse. This is consistent with the fact that cross-sectional MTV–TLA pairs are highly informative about emergent scaling behaviour, but are less able to uniquely resolve the internal dynamical decomposition of that behaviour into growth, inhibition, and saturation mechanisms.

**Figure 6.**
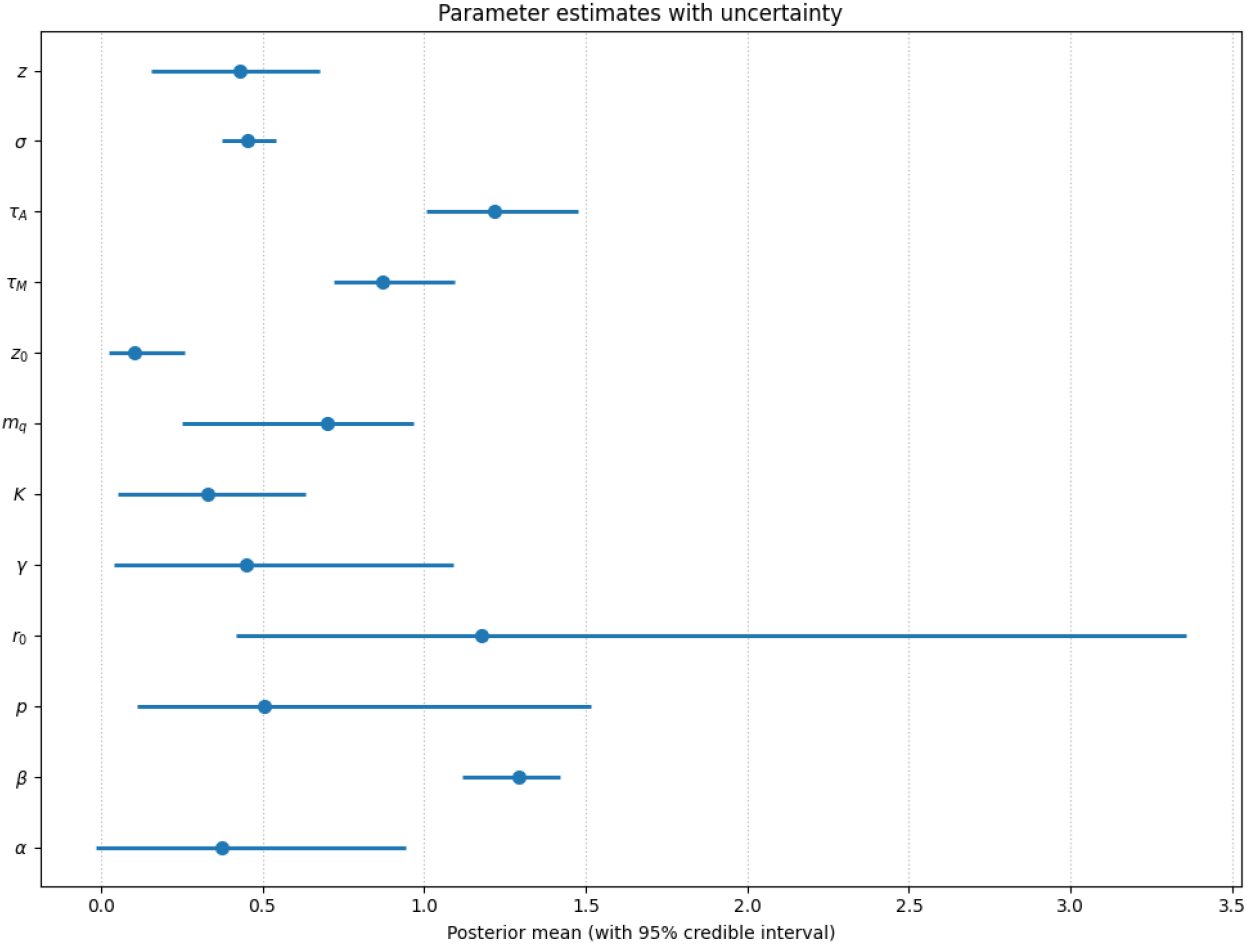
Posterior means with 95% credible intervals for the principal model parameters. The figure shows heterogeneous posterior concentration under cross-sectional observations: the effective scaling and observation-layer parameters are more tightly constrained, whereas several mechanistic parameters remain only weakly identified.

From a mechanistic perspective, the posterior intervals for parameters such as *r*_0_, *p*, and *γ* suggest that the model accommodates a range of biologically plausible PDE configurations that remain consistent with the observed cross-sectional data. In contrast, the tighter posterior concentration for *β* shows that the effective superlinear scaling law is robustly identified, even while the underlying mechanistic contributors retain some flexibility. The parameters *K, m*_*q*_, and *z* appear to be moderately informed, indicating that the available data already contain meaningful signal about saturation effects and latent-state structure, while also leaving room for these components to be sharpened further through richer data sources or complementary model constraints.

The posterior for the intercept *α* supports a positive baseline shift in the log-scale MTV–TLA relation, while the relatively concentrated posterior for *σ* indicates that the residual dispersion around this scaling law is learned with reasonable stability. This is compatible with a setting in which MTV explains an important share of the variation in TLA on the log scale, while meaningful heterogeneity remains due to tumour phenotype differences, microenvironmental effects, and imaging or segmentation variability.

Overall, the posterior structure suggests that the present cross-sectional data are well suited for learning effective scaling behaviour and observation-level variability, but are less informative about the full mechanistic decomposition of tumour progression. For stronger mechanistic identification, richer data sources, such as longitudinal imaging, lesion-level trajectories, or complementary biomarkers, would be needed to disentangle proliferation, decay, and saturation effects more reliably.

## 6 Discussion and outlook

A main contribution of the present work is the introduction of a *novel* nonlocal FKPP–Kawarada-type tumour-growth model in which proliferation is regulated by a convolution-based neighbourhood signal and undergoes singular acceleration as this nonlocal burden approaches a critical threshold. This modelling step is motivated by the experimental evidence in [1] that tumour activity can exhibit sustained superlinear scaling and an associated *explosive* escalation that is not rigorously captured by classical local logistic/FKPP dynamics. By incorporating nonlocal feedback together with a Kawarada-type singularity, our model provides a concrete dynamical mechanism capable of reproducing such explosive regimes while keeping the tumour density itself bounded, thereby offering a mesoscopic PDE interpretation of the behaviour observed in [1].

Building on this mechanistic perspective, we have analyzed the resulting singular nonlocal logistic dynamics under homogeneous Neumann boundary conditions, showing that convolution-based feedback with a power-law singularity induces finite-time quenching. In particular, spatially homogeneous states reduce the PDE to a scalar nonlocal Kawarada-type ODE for the mean density, enabling explicit characterization of the quenching time and its rate. Beyond the homogeneous setting, we established a spectral stability theory in which perturbations decompose into Neumann Laplacian eigenmodes, and the stabilizing or destabilizing influence of nonlocality is governed by the interaction kernel through its (Neumann-adapted) transform. This yields a transparent connection between spatial scales, kernel structure, and the onset of patterning or stabilization as the system approaches the critical threshold. The singularity exponent *p* emerges as a key control parameter: increasing *p* effectively acts as a tunable amplifier of nonlocal feedback, sharpening spatial gradients and accelerating the approach to quenching, while also tightening stability margins by magnifying the response to convolution-driven fluctuations.

From a mathematical viewpoint, a central contribution of this work is the integration of the deterministic PDE analysis with a Bayesian inference layer.Treating the kernel parameters and the singularity exponent as uncertain quantities, and calibrating them to tumour-growth observations, allows uncertainty to be propagated from data and modelling assumptions to the primary dynamical outputs. In this way, quenching time and quenching rate become not single predictions but posterior distributions, and spectral stability becomes a probabilistic statement about which spatial modes are likely to be amplified given uncertainty in the nonlocal coupling. This perspective is particularly valuable in explosive growth regimes, where small parameter perturbations can produce large changes in predicted escalation times. The Bayesian setting also supports principled comparison of competing kernel families and regularization choices, and it enables posterior predictive checks to identify systematic discrepancies between the quenching model and observed dynamics.

### Limitations of the nonlocal model

While the proposed nonlocal Kawarada-type formulation provides a useful mechanistic framework for capturing accelerated tumour growth, there are several aspects that warrant caution when interpreting results or transferring the model to specific datasets. First, the power-law singularity offers an idealised representation of strongly accelerated proliferation. Although this mechanism is convenient for analysing explosive regimes, it may not fully reflect additional biological processes that can moderate growth in practice (e.g., nutrient limitation, hypoxia, immune response, or treatment effects). In particular, the threshold-driven quenching structure is encoded directly in the reaction law; if tumour progression approaches such regimes more gradually, the model may accentuate the sharpness of the acceleration near the inferred threshold.

Second, the interaction kernel provides a flexible way to represent nonlocal coupling, but in many applications its detailed form may be only partially identifiable from available measurements. With coarse or primarily cross-sectional observations, different kernel shapes (and compensating choices of the exponent *p*) can lead to similar fits, so posterior inferences may depend on modelling choices and prior regularisation. Even when spatial information is available, kernel estimation can be sensitive to discretisation, boundary geometry, and the chosen observation operator. This suggests the value of careful prior specification, model comparison, and sensitivity analysis, and it highlights the potential benefits of richer data modalities when feasible.

Finally, the present formulation assumes an isotropic, time-invariant kernel and homogeneous tissue properties. Real tumour microenvironments can exhibit heterogeneity and anisotropy, including spatially varying motility, directed migration, evolving vasculature, and interaction ranges that change over time. Extending the framework to allow spatially varying or adaptive kernels would broaden biological realism, but would also introduce additional parameters and computational demands.

### Outlook

Several directions follow naturally from the present study. On the modelling side, it is of interest to incorporate additional biology, such as heterogeneous diffusion, mechanochemical coupling, immune interactions, or therapy terms, and to study how these mechanisms interact with nonlocal quenching. On the analytical side, extensions to non-symmetric kernels and more general nonlocal operators could broaden applicability while preserving the spectral interpretability established here. From the Bayesian perspective, hierarchical and partially pooled formulations could leverage multi-patient datasets to separate population-level interaction structure from individual variability, while experimental design questions could be posed to determine which data types (spatial snapshots, longitudinal imaging, biomarkers) most effectively reduce uncertainty in quenching-time predictions. Together, these developments would move the nonlocal Kawarada framework toward a more comprehensive and testable model class for explosive tumour dynamics.

